# Systemic immune dysfunction in cancer patients driven by IL6 and IL8 induction of an inhibitory receptor module in peripheral CD8^+^ T cells

**DOI:** 10.1101/2020.05.06.081471

**Authors:** Ashwin Somasundaram, Anthony R. Cillo, Caleb Lampenfeld, Lauren Oliveri, Maria A. Velez, Sonali Joyce, Michael J. Calderon, Rebekah Dadey, Dhivyaa Rajasundaram, Daniel P. Normolle, Simon C. Watkins, James G. Herman, John M. Kirkwood, Evan J. Lipson, Robert L. Ferris, Tullia C. Bruno, Dario A.A. Vignali

## Abstract

Many cancer patients do not develop a durable response to the current standard of care immunotherapies despite substantial advances in targeting immune inhibitory receptors^1-5^. A potential compounding issue, which may serve as an unappreciated, dominant resistance mechanism, is an inherent systemic immune dysfunction that is often associated with advanced cancer^6-12^. Minimal response to inhibitory receptor (IR) blockade therapy and increased disease burden have been associated with peripheral CD8^+^ T cell dysfunction, characterized by suboptimal T cell proliferation and chronic expression of IRs (eg. Programmed Death 1 [PD1] and Lymphocyte Activation Gene 3 [LAG3])^13, 14^. Here, we demonstrate that up to a third of cancer patients express robust intracellular LAG3 (LAG3^IC^), but not surface LAG3 (LAG3^SUR^), in peripheral CD8^+^ T cells compared to CD4^+^ T cells and regulatory T cells (T_regs_). LAG3^IC^ is associated with: (i) expression of a LAG3^IC^-dominant IR module that includes PD1^IC^, NRP1^IC^, CD39^IC^, and TIGIT^IC^; (ii) decreased CD8^+^ but not CD4^+^ T cell function that can be reversed by anti-LAG3 (and/or anti-PD1), despite limited constitutive surface IR expression; and (iii) poor disease prognosis. Systemic immune dysfunction is restricted to CD8^+^ T cells, including a high percentage of peripheral naïve CD8^+^ T cells, indicating a TCR-independent mechanism that is driven by the cytokine IL6 and the chemokine IL8. Thus, the combination of an increased LAG3-dominant IR module and elevated systemic IL6 and/or IL8 may serve as predictive biomarkers and increase the possibility that cancer patients will benefit from therapeutic combinations targeting these systemic cytokines in the setting of PD1 and/or LAG3 blockade.

---

PD1/PDL1 blockade has revolutionized therapeutic options for solid tumors^1, 3-5, 15, 16^, however, only 20-30% of patients will respond to PD1 blockade, emphasizing the need to identify underlying mechanisms of immune resistance^17^. Another important and often underappreciated fact is that many cancer patients have reactivation of chronic viral infections, lose their response to vaccinations and succumb to secondary infections without tumor progression^7-12^. Expression of additional inhibitory receptors (IRs) on peripheral immune cells has been associated with immune resistance across multiple tumor types^13, 18, 19^. The extent to which these observations underlie a dominant mechanism of immune resistance to immunotherapy and whether this is connected to general immune dysfunction in cancer patients remains unknown. LAG3 is an IR that has been associated with a dysfunctional CD8^+^ T cell phenotype in the periphery of mouse models of cancer and metastatic melanoma patients^13, 18-21^. Although surface LAG3 (LAG3^SUR^) expression on peripheral blood T cells can be limited, we speculated that LAG3 expression may be more impactful than previously appreciated due to: (i) rapid cell surface shedding of LAG3 via the metalloproteases ADAM10 and ADAM17, and (ii) prevalent intracellular LAG3 storage in T cells^22-25^. Thus, we evaluated the dynamics of surface and intracellular LAG3 expression, the expression of other IRs, and the potential systemic factors that might drive systemic immune dysfunction in cancer.

We initially analyzed single-cell RNA sequencing (scRNAseq) data from 26 HNSCC patient peripheral blood leukocytes (PBL) and 6 healthy donors (HD) PBL (**Fig. 1a**)^26^. Interestingly, *LAG3* was expressed in a high percentage of T cells and appeared to be largely restricted to HNSCC patient CD8^+^ T cells over other cell populations, including patient CD4^+^ T effector cells and T_regs_, and HD CD8^+^ T cells (**Fig. 1a, Extended Data Fig. 1a**). A similar pattern of expression has also been reported in NSCLC patients^27^. Further analysis suggested that *PDCD1* (encoding PD1) exhibited a similar expression pattern, albeit to a lesser extent. These transcriptomic data suggest that *LAG3* and *PDCD1* are expressed at increased levels in peripheral CD8^+^ T cells of some cancer patients, an observation that had not previously been appreciated (**Fig. 1a, Extended Data Fig. 1a-c**).

**Figure 1:**
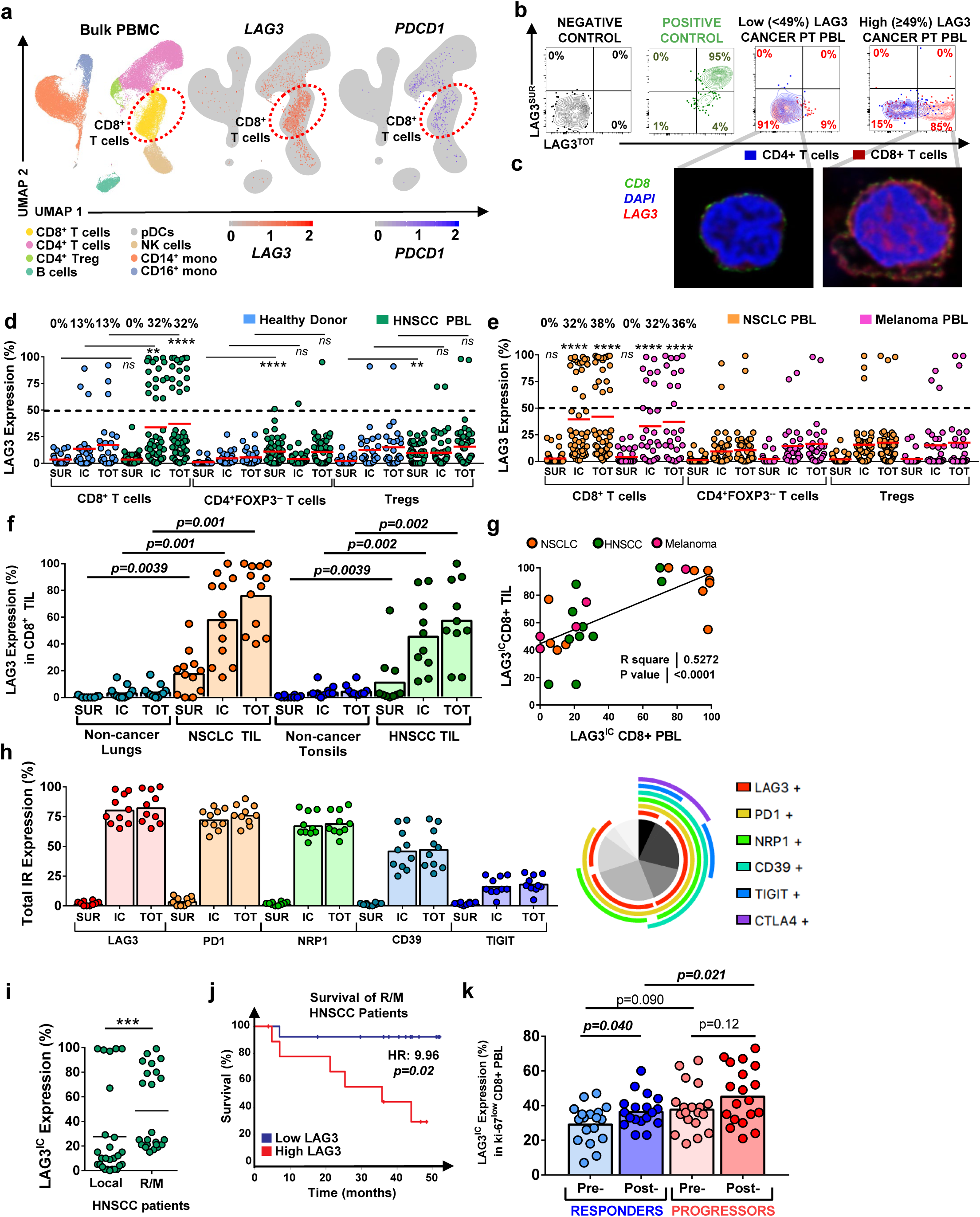
A LAG3-dominant IR module in peripheral CD8^+^ T cells of cancer patients is associated with worse clinical outcomes and immunotherapeutic resistance. **a**, tSNE plots of unsupervised clustering of 26 HNSCC patients and 6 HD sample cells by cell type with *LAG3* (red) and *PDCD1* (blue) gene expression across cell types. **b**, Representative flow cytometric plots of CD8^+^ T cells from a healthy donor (black), CD8^+^ T cells activated with anti-CD3 (0.5μg/mL) and anti-CD28 (1μg/mL) as a positive control (green), CD4^+^ (blue) and CD8^+^ (red) T cells from a patient with cancer and with low (<49%) total LAG3 on their CD8^+^ T cells and from a patient with cancer and high (≥49%) total LAG3 on their CD8^+^ T cells. **c**, STED microscopy of a CD8^+^ T cell from an HNSCC patient with low intracellular LAG3, and a CD8^+^ T cell from an HNSCC patient with high intracellular LAG3. The markers used are CD8 (green), DAPI for nuclear staining (blue), LAG3 (red). **d**, Surface, intracellular, and total LAG3 expression by flow cytometry in HNSCC (green; n=50) and HD PBL (blue; n=26) across T cell subsets including: CD8^+^ T cells, CD4^+^ T cells, and Tregs. A mean cutoff of greater than or equal to 49% LAG3^TOT^ expression is represented by the black dashed line. Percent of patients above the mean cutoff is noted above the statistics. *P <0.05, ***P <0.001, ****P< 0.0001, *ns* =not significant or P >0.05, Wilcoxon matched-pairs test. **e**, Surface, intracellular, and total LAG3 expression by flow cytometry in NSCLC (orange; n=50) and metastatic melanoma PBL (pink; n=28) across T cell subsets including: CD8^+^ T cells, CD4^+^ T cells, and Tregs. LAG3 expression among each T cell subset is compared to its matched subset from HD PBL (**Fig 1d**). The percentage of patients above the mean cutoff of 49% (black dashed line) is noted as a percent above the statistical symbols. *P <0.05, ***P <0.001, ****P< 0.0001, *ns* =not significant or P >0.05, Wilcoxon matched-pairs test. **f**, Surface, intracellular, and total LAG3 expression by flow cytometry in TIL from patients. compared to tissue controls. Healthy donor lung tissue (n=11) compared to NSCLC TIL (n=12), and healthy donor tonsil (n=10) compared to HNSCC TIL (n=10). Wilcoxon matched-pairs test. **g**, Spearman’s correlation of matched PBL and TIL from high and low intracellular LAG3 patients (NSCLC, n=12; HNSCC, n=10; and metastatic melanoma n=5). **h**, Surface, intracellular, and total LAG3 expression by flow cytometry of a total IR module consisting of LAG3, PD1, NRP1, CD39, TIGIT from only high intracellular LAG3 patients with NSCLC. Matched SPICE plot depicting the co-expression of IRs within these selected patients. **i**, Comparison of LAG3^IC^ expression from local vs. recurrent/metastatic (R/M) HNSCC patients (**Fig. 1d**). *P <0.05, ***P <0.001, ****P< 0.0001, ns =not significant or P >0.05, Wilcoxon matched-pairs test. **j**, Survival of patients with low (<49%) LAG3^IC^ CD8^+^ PBL (blue) versus survival of patients with high (≥49%) LAG3^IC^ CD8^+^ PBL from the R/M subgroup in **Fig 1i**. Hazard ratio (HR) of 9.96 (95% confidence interval: 1.176 to 177; p=0.02) after multivariate analysis including future treatments (systemic therapy versus local therapy alone), gender, tobacco use, and HPV status. *P <0.05, ***P <0.001, ****P< 0.0001, *ns* =not significant or P >0.05, Wilcoxon matched-pairs test. **k**, Comparison of LAG3^IC^ expression in ki-67^low^ CD8^+^ T cells by flow cytometry from both responder (blue; n=18) and progressor (red; n=19) patients with metastatic melanoma and other skin cancers before checkpoint blockade and 12 weeks on checkpoint blockade at the time of their initial scans (4 cycles of therapy). Wilcoxon matched-pairs test.

Despite increased transcription, LAG3^SUR^ expression on patient peripheral T cells by flow cytometric analysis was minimal across all immune cell subpopulations compared to CD8^+^ T cells (**Fig. 1b-e**). Our previous studies with mouse T cells suggest that a substantial amount of LAG3 can be stored intracellularly, awaiting rapid trafficking to the cell surface following TCR stimulation^23^. Intracellular, but not surface, LAG3 has been reported as elevated in cancer patient peripheral blood T cells compared to healthy donors^25^. Using two non-competing antibodies, we analyzed LAG3^SUR^ and intracellular LAG3 (LAG3^IC^; determined by subtracting LAG3^SUR^ from LAG3^TOT^) by multi-parameter flow cytometry and by stimulated emission depletion (STED) microscopy (**Fig. 1b, c**). Approximately 30% of patients with HNSCC, NSCLC, and metastatic melanoma exhibited a striking bimodal distribution of LAG3^TOT^ and LAG3^IC^ protein expression; ‘high’ LAG3^TOT^/LAG3^IC^ defined as 49-100% expression and ‘low LAG3^TOT^/LAG3^IC^ defined as 0-49% expression. (**Fig. 1d, e**). This was a unique feature of CD8^+^ T cells as this was not observed with other patient T cell populations (eg. CD4^+^Foxp3^−^ T cells and T_regs_) and HD T cells. While these findings were observed in the peripheral blood, high LAG3^IC^ expression on matched tumor-infiltrating lymphocytes (TIL) also had a positive correlation with high LAG3^IC^ expression in peripheral CD8^+^ T cells, suggesting that these systemic findings may inform the state of the tumor microenvironment (**Fig. 1f, g**). Interestingly, patients with high LAG3^IC^ in their peripheral CD8^+^ T cells also exhibited co-expression of other IRs, notably PD1^IC^, NRP1^IC^, CD39^IC,^ and TIGIT^IC^ (**Fig. 1h, Extended Data Fig. 1d, e, 2**), ultimately creating a LAG3-dominant module of IR expression.

To further evaluate the state of chromatin modeling at the site of these IR genes, genes of T cell naivety, and genes of T cell activation^28^, we performed ATAC-seq on CD8^+^ T cells from NSCLC PBL from both patients with low and high LAG3^IC^-dominant IR expression and found that relative to areas of known closed chromatin as a negative control, the IR associated areas of chromatin were open (**Extended Data Fig. 3**). However, there were no differences in chromatin accessibility between the low vs. high LAG3^+^ CD8^+^ T cells, suggesting that regulation of expression may happen downstream of chromatin remodeling.

A blinded, retrospective analysis of 50 HNSCC patient PBL samples indicated worse disease burden i.e. recurrent/metastatic disease (R/M), and worse overall survival in patients with high LAG3^IC^ in their peripheral CD8^+^ T cells (**Fig. 1i, j, Supplementary Table 1**). Taken together, these data suggest that a LAG3-dominant IR module in systemic CD8^+^ T cells may contribute to a reduction of productive immunity in cancer patients. Further, these data also suggest that systemic IR expression should be considered as a biomarker of increased disease burden.

To better interrogate if LAG3^IC^ may also predict response to immunotherapy, we evaluated a cohort of 37 patients with skin cancer malignancies that received standard of care immunotherapies (**Supplementary Table 2**). Interestingly, we found that LAG3^IC^ was increased more significantly in the Ki-67^low^ CD8^+^ T cells in the post timepoints of progressors vs. responders, ultimately suggesting that this might serve as a biomarker for immunotherapeutic resistance (**Fig. 1k, Extended Data Fig. 4**).

We next interrogated LAG3 kinetics as intracellular storage of LAG3 does not allow it to engage with its ligand, ultimately leading to two key questions: (a) does LAG3^IC^ shuttle to the cell surface, and (b) does LAG3^IC^ have a functional consequence on CD8^+^ T cells? LAG3 function is dependent on cell surface expression and ligand interaction^20^, however, the majority of our findings have been intracellular LAG3. We first expanded our STED experiments to evaluate intracellular localization of LAG3 expression via co-expression with key intracellular trafficking proteins (Cathepsin D, LAMP2, Rab27a, Rab5a, Rab11a, EEA1). These data suggested that intracellular LAG3 co-localizes with vesicles containing Cathepsin D and/or Early endosome antigen 1 (EEA1), which raises the possibility that LAG3 is cycling via endosomes (**Extended Data Fig. 5)**^29-33^. While this finding confirms intracellular compartmentalization, it doesn’t provide sufficient insight into the surface trafficking of the protein. Thus, we first evaluated LAG3^SUR^ expression over time on stimulated CD8^+^ T cells from NSCLC patients with high and low LAG3^IC^ compared to HD. We found that CD8^+^ T cells from patients with high LAG3^IC^ had higher LAG3^SUR^ expression at 24h and 48h following *in vitro* TCR stimulation compared to patients with low LAG3^IC^ or HD (**Fig. 2a**). We also found that LAG3^IC^ and LAG3^TOT^ expression in CD8^+^ T cells strongly correlated with soluble LAG3 (sLAG3) release into the media after a 24h *in vitro* culture with TCR stimulation (co-culture with plate-bound anti-CD3 and anti-CD28) (**Fig. 2b**). This suggests that LAG3 is not simply retained intracellularly, but rather, is dynamically translocated to the cell surface to be subsequently shed.

**Figure 2:**
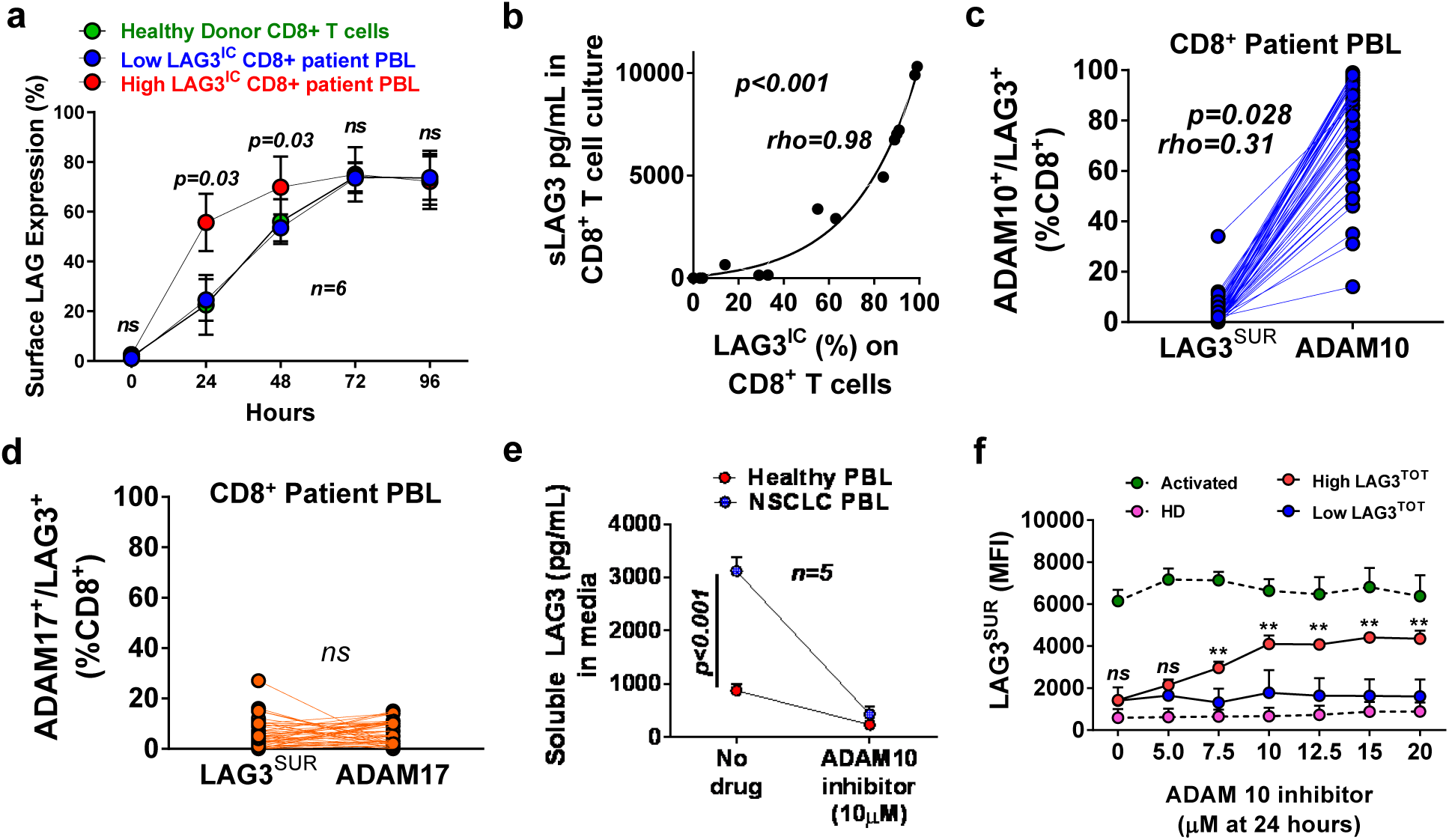
LAG3 is stored intracellularly in CD8+ T cells and traffics to the cell surface with rapid shedding via ADAM 10. **a**, Surface LAG3 expression over time in CD8^+^ T cells from healthy donors (n=6), from low LAG3 NSCLC patients (n=6), and high LAG3 NSCLC patients (n=6), during stimulation with anti-CD3 (0.5μg/mL) and matched 1:1 APCs. Wilcoxon matched-pairs test. **b**, Spearman’s correlation of soluble LAG3 in media culture and LAG3^IC^ expression on CD8^+^ T cells from patients during 24 hours of TCR stimulation with plate-bound anti-CD3 and anti-CD28 (n=14). **c**, Spearman’s correlation of matched ADAM10 and LAG3^SUR^ expression on CD8^+^ T cells from HNSCC patients (n=49). **d**, Spearman’s correlation of matched ADAM17 and LAG3^SUR^ expression on CD8^+^ T cells from HNSCC patients (n=49). **e**, Soluble LAG3 in culture media of CD8^+^ T cells from NSCLC patients (n=5) compared with CD8^+^ T cells from healthy donors (n=5), with and without the presence of an ADAM10 inhibitor (10μM). **f**, LAG3^SUR^ expression by MFI on CD8^+^ T cells from healthy donors (negative control; n=5), activated CD8^+^ T cells (positive control; n=5), CD8^+^ T cells from patients with low LAG3^IC^ (n=5), and high LAG3^IC^ (n=5) at increasing dose titrations of an ADAM10 inhibitor without the presence of TCR stimulation.

We hypothesized that the presence of LAG3^IC^ and the absence of LAG3^SUR^ may be due to constitutive shedding by ADAM family metalloproteinases with the release of sLAG3^22^. Consistent with this hypothesis, we found that most peripheral CD8^+^ T cells exhibited an inverse correlation between LAG3^SUR^ and ADAM10, but not ADAM17, surface expression with the predominant pattern being low LAG3^SUR^ and high ADAM10 expression in the CD8^+^ T cells (**Fig. 2c, d**)^22^. Indeed, LAG3^SUR^ expression increased and sLAG3 in media decreased after *in vitro* treatment with an ADAM10 inhibitor (**Fig. 2e, f**). These data imply that LAG3 does reach the cell surface but is rapidly shed by ADAM10 explaining the lack of detectable cell surface expression by flow cytometry. Thus, LAG3 protein trafficking to the cell surface does occur in the presence of TCR stimulation, but only for a limited time before being shed by ADAM10. These data suggest that LAG3^IC^ could have a functional consequence in peripheral CD8^+^ T cells from patients with cancer.

We interrogated the functional consequence of high LAG3^IC^ expression on peripheral CD8^+^ T cells in a cohort of NSCLC patient samples. CD8^+^ T cells from patients with high LAG3^IC^ expression demonstrated decreased cytokine production (TNFα, IL2, IFNγ) upon stimulation with PMA and ionomycin compared to CD8^+^ T cells from patients with low LAG3^IC^ expression or HD, consistent with reported findings in murine models (**Fig. 3a, b**)^34^. CD8^+^ T cells with high LAG3^IC^ from patients with either NSCLC or metastatic melanoma also exhibited reduced T cell proliferation (**Fig. 3c, d**). Strikingly, CD8^+^ T cell proliferation could be restored with the addition of either anti-PD1, anti-LAG3, or the combination (**Fig. 3e-g**). Taken together, despite minimal detectable LAG3^SUR^ expression by flow cytometry due to rapid ADAM-mediated shedding (**Fig. 2a-c**), IR blockade can still reverse T cell dysfunction *in vitro*. These data raise the possibility that IR blockade may not only reverse intratumoral T cell dysfunction but may also relieve systemic T cell immune dysfunction despite rapid shedding of LAG3 from the surface of CD8^+^ T cells.

**Figure 3:**
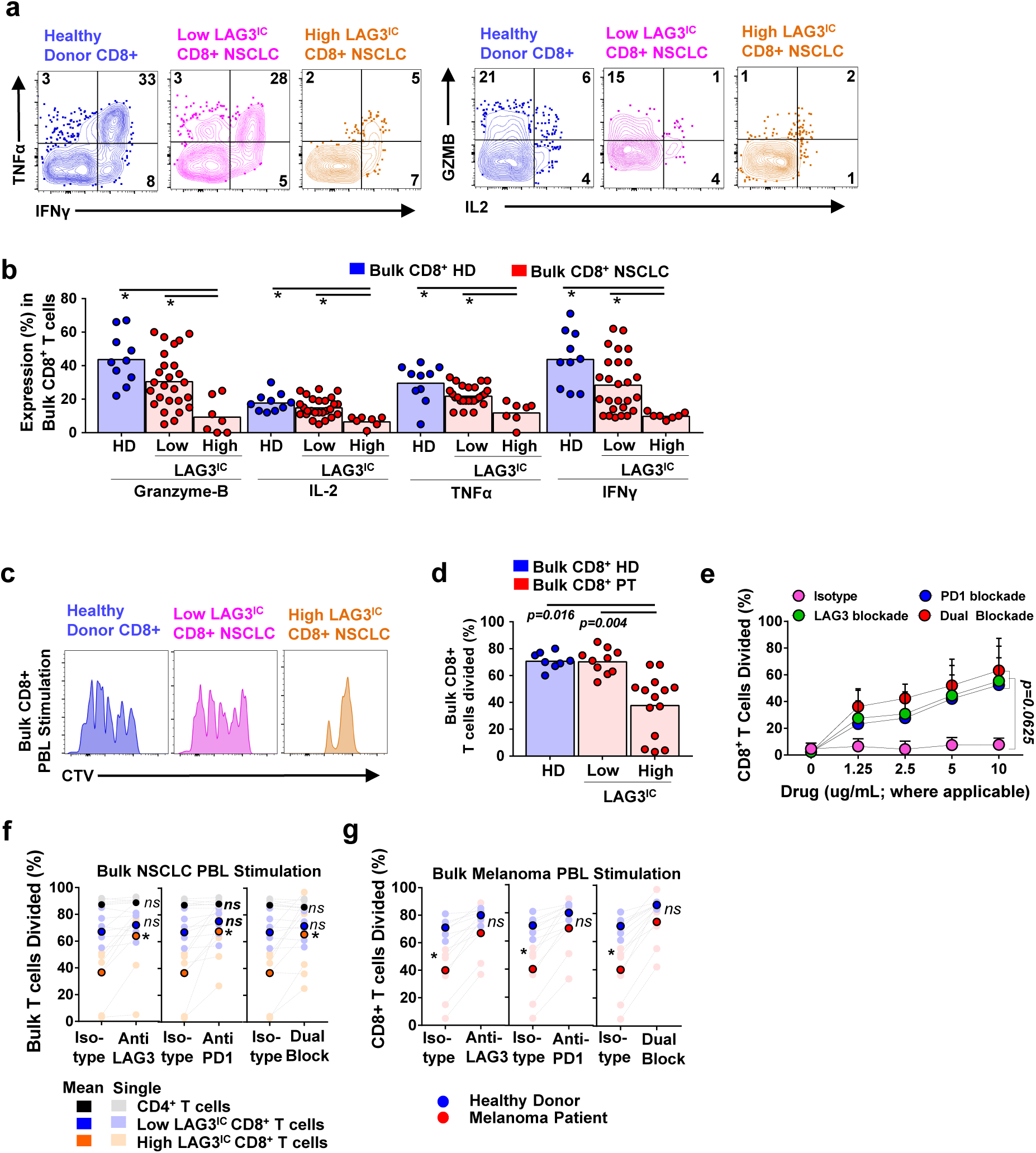
Peripheral CD8^+^ T cells with an elevated LAG3-dominant IR module have reduced function that can be partially rescued by anti-LAG3 or anti-PD1. **a**, Representative flow cytometric plots of TNFα, IFN γ, granzyme-B (GZMB), and IL2 expression in CD8^+^ peripheral T cells from a healthy donor (blue), in CD8^+^ PBL from a patient with NSCLC and low LAG3^IC^ expression (pink), and in CD8^+^ PBL from a patient with NSCLC and high LAG3^IC^ expression (orange) after stimulation with PMA and ionomycin for 4-6 hours. **b**, TNFα, IL2, IFNγ and granzyme B production evaluated by flow cytometry after 4-6 hour PMA and ionomycin stimulation in bulk CD8^+^ T cells with elevated LAG3^IC^/PD1^IC^ expression from NSCLC patients (n=7) compared to CD8^+^ T cells with low LAG3^IC^/PD1^IC^ (n=26) and CD8^+^ T cells from HD (n=10). Statistical significance determined by Wilcoxon signed-rank test (ns is P>0.05, * is P≤0.05, ** is P≤0.01, *** is P≤0.001, **** is P≤0.0001). **c**, Representative flow cytometric histogram plot of CTV-labelled proliferation in CD8^+^ PBL from a patient with NSCLC and high LAG3^IC^ expression (blue), and CD8^+^ PBL from a patient with NSCLC and low LAG3^IC^ expression (red). **d**, Proliferation of CTV-labelled CD8^+^ bulk PBL by flow cytometry from health donors (HD; n=8), patients with NSCLC and low LAG3^IC^ (n=5), patients with metastatic melanoma and low LAG3^IC^ (n=5), patients with NSCLC and high LAG3^IC^ (n=7), or patients with metastatic melanoma and high LAG3^IC^ (n=7). Proliferation assay was conducted over 96 hours with CD8+ T cells incubated with matched APCs, and anti-CD3 (0.5μg/mL). (Statistical significance determined by Wilcoxon signed-rank test (ns is P>0.05, * is P≤0.05, ** is P≤0.01, *** is P≤0.001, **** is P≤0.0001). **e**, Cells from NSCLC PBL that contain the high LAG3^IC^ CD8^+^ T cell population were stimulated with anti-CD3 (0.5μg/mL) and matched 1:1 APCs at a 96-hour timepoint in the presence of anti-LAG3 and/or anti-PD1 and T cell division was assessed by CTV-labelling. Drug or isotype was titrated at concentrations of 10µg/mL, 5µg/mL, 2.5µg/mL, 1.25µg/mL, and 0µg/mL. *P <0.05, ***P <0.001, ****P< 0.0001, ns is P>0.05, Wilcoxon matched-pairs test. Error bars, s.d. **f**,, Bulk CD4^+^ T cells (n=5) and CD8^+^ T cells (n=12) from NSCLC patients with high (n=7) and low LAG3^IC^ (n=5) expression, were stimulated with anti-CD3 (0.5μg/mL) and matched 1:1 APCs at a 96 hour time point in the presence of anti-LAG3 (10μg/mL) and/or anti-PD1 (10μg/mL). Correlations were performed with Spearman’s nonlinear correlation. Mean values of proliferation noted in darker colors and individual sample proliferation noted in lighter colors. Statistical significance determined by Wilcoxon signed-rank test (ns is P>0.05, * is P≤0.05, ** is P≤0.01, *** is P≤0.001, **** is P≤0.0001) comparing the change in proliferation after the addition of checkpoint blockade. **g**, CD8^+^ T cells from healthy donors (n=5), and patients with metastatic melanoma and high LAG3^IC^ expression (n=7) were stimulated with anti-CD3 (0.5μg/mL) and matched 1:1 APCs at a 96-hour timepoint in the presence of anti-LAG3 (10μg/mL) and/or anti-PD1 (10μg/mL). Correlations were performed with Spearman’s nonlinear correlation. Mean values of proliferation noted in darker colors and individual sample proliferation noted in lighter colors. Statistical significance determined by Wilcoxon signed-rank test (ns is P>0.05, * is P≤0.05, ** is P≤0.01, *** is P≤0.001, **** is P≤0.0001) comparing the change in proliferation after the addition of checkpoint blockade.

One of the most surprising aspects of these data is that in some patients essentially all their peripheral CD8^+^ T cells display high expression of this IR module, exemplified by LAG3^IC^ expression. Extensive immunophenotyping of patient peripheral CD8^+^ T cells revealed that this LAG3-dominant IR module was also expressed by naïve CD8^+^ T cells along with other T cell subsets, such as effector memory CD8^+^ T cells and terminally differentiated (T_EMRA_) CD8^+^ T cells (**Extended Data Fig. 6a, b**)^35^. One immediate conclusion is that these CD8^+^ T cells are not bona fide naïve CD8^+^ T cells, and perhaps have endured an activation or proliferation event while still exhibiting a canonical naïve T cell phenotype (CD45RA^+^, CCR7^+^, CD62L^+^, CD127^+^, CD95^−^, CD69^−^, HLA-DR^−^)^36^. We utilized analysis of T cell receptor excision circles (TREC) to evaluate their ‘proliferative history’ and state of naivety. During TCR rearrangement, TREC is generated and their prevalence among T cells has been used as a marker of their naive status and lack of TCR-driven proliferation^37-40^. TREC concentrations were similar between purified naive CD8^+^ T cells from patients with or without an elevated LAG3-dominant IR module and purified naïve HD CD8^+^ T cells (**Extended Data Fig. 6c)**. These data suggest that patients with high LAG3^IC^ expression in their naïve CD8^+^ T cells are not expressing the LAG3-dominant IR module due to a proliferative event that might otherwise induce IR expression^37^. We also queried the functional consequence of the LAG3-dominant IR module concerning naïve CD8^+^ T cell function and found that cytokine production and proliferation were also affected in this subset and this T cell dysfunction could be rescued with checkpoint blockade (**Extended Data Fig. 7a-d**). Taken together, these data suggest that a cell-extrinsic stimulus is driving the expression of a LAG3-dominant IR module on bona fide naïve CD8^+^ T cells that would presumably not have been exposed to a cognate TCR stimulation or activation event.

Given the influx of inflammatory factors into the blood of cancer patients, we first asked if systemic cytokines in plasma could provide an extrinsic stimulus driving IR expression in naïve CD8^+^ T cells. Remarkably, we found that plasma from NSCLC patients with high LAG3^IC^, but not NSCLC patients with low LAG3^IC^ or heavy smoker ‘healthy’ control patients, could induce LAG3IC and PD1^IC^ expression in HD naïve CD8^+^ T cells without TCR stimulation (**Fig 4a, b**). We further corroborated these data with plasma from patients with metastatic melanoma or HNSCC who expressed high vs. low LAG3^IC^ (**Extended Data Fig. 8a, b**) for an array of cytokines while simultaneously testing if recombinant versions of these cytokines could induce LAG3^IC^ and PD1^IC^ in HD CD8^+^ T cells. Interestingly, six cytokines were both elevated in patient plasma and could induce LAG3^IC^ and PD1^IC^ in naïve CD8^+^ T cells in the absence of TCR stimulation: IL6, IL8, IL9, IL10, IL15 and IL21 (**Fig. 4c, Extended Data Fig. 8c-e, Supplementary Table 3**). Further, all of these cytokines had a statistically significant correlation with LAG3^IC^ and PD1^IC^ expression in CD8^+^ T cells (**Fig. 4c, Supplementary Table 3**). We further evaluated the induction of the other IRs in this module in the presence of IL6 and IL8 and noted a trend towards induction while not statistically significant (**Extended Data Fig. 8e**).

**Figure 4:**
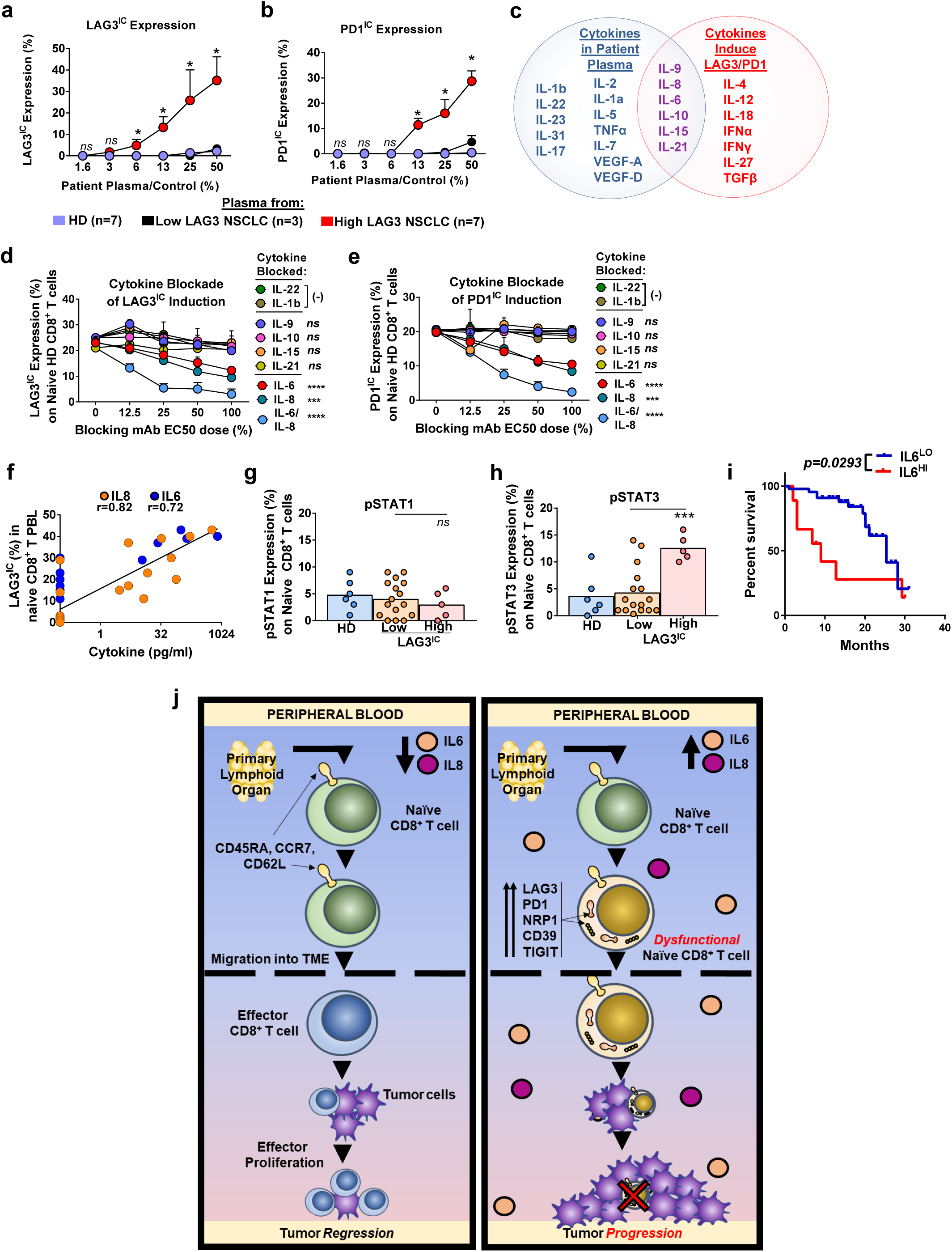
LAG3-dominant IR module in peripheral, naïve CD8^+^ T cells is driven by IL6 and IL8. **a, b**, Naïve (CD45RA^+^CCR7^+^CD62L^+^) CD8^+^ T Cells from healthy donors were incubated with IL2 and IL7 and plasma from patients with high LAG3^IC^/PD1^IC^ on naïve CD8^+^ T cells (n=7), plasma from patients with low LAG3^IC^/PD1^IC^ on naïve CD8^+^ T cells (n=3), and plasma from healthy donors (n=7) for 60 hours and analyzed for LAG3^IC^ or PD1^IC^ expression.) *P <0.05, ***P <0.001, ****P< 0.0001, *ns* is P>0.05, Wilcoxon matched-pairs test. Error bars, s.d. **c**, Venn Diagram of all cytokines with any similar correlation between their plasma concentrations and LAG3^IC^/PD1^IC^ expression on their naïve CD8^+^ T cells (*blue*) followed by their ability to induce LAG3/PD1 expression in an *in vitro* incubation assay (*red*). Cytokines that are both correlated and can induce LAG^IC^/PD1^IC^ expression are noted in purple. For full Spearman’s correlation rho values between plasma concentrations of cytokines and total/intracellular LAG3 or PD1 expression on their naïve CD8^+^ T cells *see Extended table 2.* **d, e**, Naïve CD8^+^ HD T cells incubated with plasma from High LAG3^IC^/PD1^IC^ NSCLC patients in a similar manner to **Fig 4. a, b**. This plasma was known by multiplex to have elevated concentrations of all the cytokines of interest that correlated with LAG3^IC^/PD1^IC^. Incubation also included blockade of specific cytokines (anti-IL6 n=6; anti-IL8 n=7; anti-IL9 n=4; anti-IL10 n=3; anti-IL15 n=3; anti-IL22 n=4; anti-IL1β n=4, anti-IL21 n=5; anti-IL6/IL8 n=5) in an effort to rescue plasma-induced LAG3^IC^/PD1^IC^ expression. Concentrations of each blocking antibody vary, and the x-axis is noted as the percent of the reported EC_50_ blocking dose. IL22 and IL1β concentration shown to be elevated by multiplex noted in *Extended Table 2* and correlated with LAG3^IC^/PD1^IC^ but **not** noted to be able to induce LAG3^IC^/PD1^IC^ in the *in vitro* induction assay, allowing blockade of IL22 and IL1β to serve as a negative control of rescue. Statistical significance determined by Wilcoxon signed-rank test (ns is P>0.05, * is P≤0.05, ** is P≤0.01, *** is P≤0.001, **** is P≤0.0001). Error bars, s.d. **f**, Spearman’s correlation of IL6 and IL8 versus naïve (CD45RA^+^, CCR7^+^. CD62L^+^) CD8^+^, LAG3^IC^ expression in NSCLC patients. Linear regression noted within data points with positive cytokine concentration. **g, h**, Cells were assessed *ex vivo.* pSTAT1 and pSTAT3 expression evaluated by flow cytometry in naïve CD8^+^ T cells from patients with elevated LAG3^IC^/PD1^IC^ expression (n=5) compared to patients with low LAG3^IC^/PD1^IC^ (n=16).Statistical significance determined by Wilcoxon signed-rank test (ns is P>0.05, * is P≤0.05, ** is P≤0.01, *** is P≤0.001, **** is P≤0.0001). **i**, Survival of advanced NSCLC patients on standard of care PD1 blockade therapy with IL6^HI^ plasma (defined as ≥ 10ng/mL) compared with advanced NSCLC patients on therapy with IL6^LO^ plasma (defined as <10ng/mL). The log-rank hazard ratio of 0.4056 (95% confidence interval: 0.1294 to 1.272; p=0.029) after multivariate analysis including only gender, tobacco use. *P <0.05, ***P <0.001, ****P< 0.0001, *ns* =not significant or P >0.05. **j**, Overall schema of cytokine inducing systemic immune dysfunction.

Next, we tested which of these six core cytokines were required to induce LAG3^IC^ and PD1^IC^ expression by assessing if neutralizing antibodies against these cytokines could block the ability of plasma from patients with high LAG3^IC^ to induce LAG3^IC^/PD1^IC^. Blockade of IL6 or IL8, but not the other cytokines, limited LAG3^IC^ and PD1^IC^ expression on HD naïve CD8^+^ T cells, with a clear additive effect with IL6/IL8 combinatorial blockade (**Fig. 4d, e, Extended Data Fig. 9a**). When plasma cytokine levels were elevated, LAG3^IC^ was increased on the naïve CD8^+^ T cells from NSCLC patients (**Fig. 4f**). We also evaluated the simultaneous effects of IL6 and IL8 with a titration of TCR stimulation as productive TCR engagement can also drive LAG3^IC^ expression. IL6 and IL8 similarly induced LAG3^IC^ expression, except for high-dose TCR stimulation (10 µg/ml), which leads to maximal expression of LAG3 (**Extended Data Fig. 9b**).

IL6 induces signal transduction via Signal Transducer and Activator of Transcription 3 (STAT3). Interestingly, elevated pSTAT3, but not pSTAT1, was observed in naive CD8^+^ T cells, but not naïve CD4^+^ T cells, in NSCLC patients with high, but not low, LAG3^IC^ (**Fig 4g, h**). NSCLC patients with high LAG3^IC^ in their CD8^+^ T cells had lower IL6 receptor (IL6R) expression and IL8 receptor (CXCR1) expression compared to CD8^+^ T cells from patients with NSCLC and low LAG3^IC^ expression (**Extended Data Fig. 9c**). Finally, evaluation of plasma IL6 from advanced NSCLC patients during standard-of-care PD1 blockade therapy revealed worse overall survival in patients with elevated plasma IL6 as previously described compared with patients with lower plasma IL6, and we observed a significant increase in the entire IR module when patients were high for systemic IL6 (**Fig 4i, Extended Data Fig. 9d**).

Taken together, these data suggest that IL6 and IL8 are elevated systemically in a subset of cancer patients and are the primary drivers of a LAG3-dominant IR module, exemplified by LAG3^IC^ and PD1^IC^, on a large proportion of CD8^+^ T cells that includes naïve T cells, but not CD4^+^ T cells and T_regs_, leading to their dysfunction and poor clinical outcome in those patients (**Fig. 4j**). It has long been known that some cancer patients have systemic immune dysfunction and worse outcomes regardless of therapy^7, 41^, but the mechanisms driving these observations were unknown. Our data suggest a novel and targetable mechanism to explain this finding that may also inform future clinical trial design. First, our study presents a cohort of biomarkers (LAG3-dominant IR module: LAG3^IC^, PD1^IC^, NRP1^IC^, CD39^IC^ & TIGIT^IC^ on a high percentage of peripheral CD8^+^ T cells, and elevated IL6 and/or IL8 in plasma) that identify patients that have a poor cancer prognosis but may respond to combinatorial therapy that targets all, or a subset, of these biomarkers. Similar findings regarding IL6 induction of LAG3 and PD1 in tumor-infiltrating CD8^+^ T cells in cancer patients has been reported^42^. Similar findings have been recently been reported for other patients with solid tumors, and combined IL6 and checkpoint blockade has shown promise in animal models^43-47^. Second, our observations highlight the necessity to fully evaluate LAG3 expression (both intracellularly and on the cell surface) in cancer immunotherapy clinical trials. Monitoring patients for any early sign of primary resistance to immunotherapy using peripheral blood CD8^+^ T cell signatures or by plasma cytokine concentrations would be both convenient and impactful for patients and clinicians as cancer patients can decline rapidly before reaching next-line therapy. Third, our findings highlight a mechanism that drives a high proportion of peripheral CD8^+^ T cells (but not peripheral CD4^+^ T cells), including naive T cells, to exhibit profound immune dysfunction as a consequence of expression of an IL6/IL8-induced LAG3-dominant IR module. IL6 and IL8 have both been reported in association with systemic inflammation of malignancy and worse outcomes of disease^48, 49^. Further prospective validation is necessary to establish if IL6/IL8 and LAG3^IC^ are resistance mechanisms to cancer therapy, especially checkpoint blockade immunotherapies, adoptive CAR-T/TIL therapies, and vaccine strategies (prophylactic and therapeutic, oncolytic, etc.)^6-9, 41, 50^. It will also be important to determine if the expression of this LAG3-dominant IR module is directly responsible for limiting the capacity of patients with advanced cancer to ward off infections that might otherwise increase morbidity or limit responsiveness to vaccination, such as the annual influenza vaccine^10-12, 51^. These findings would also impact the clinical evaluation of patients at risk for T cell dysfunction such as patients with chronic viral conditions or patients who suffer from severe autoimmunity^52, 53^. Fourth, our findings of dysfunctional naïve CD8^+^ T cells expressing IRs describes a new aspect of T cell biology that has not currently been appreciated and may be broadly applicable beyond cancer. Indeed, it has not escaped our notice that early observations in patients with COVID19 that had a poor prognosis also exhibited elevated IL6 and IR expression, suggesting our findings may have general applicability in multiple disease settings^52, 54, 55^. These findings have promising implications for combinatorial immunotherapeutic approaches targeting these cytokines and IRs, such as PD1 and LAG3, and present a set of biomarkers that may inform cancer therapy approaches.

## Acknowledgments

The authors wish to thank Jody Bonnevier at R&D Systems for providing our listed cytokines and functional antibodies. We also wish to thank BMS for providing staining and functional antibodies. The authors thank M. Meyer, B. Janjic, and A.D. Donnenberg from the HCC Flow Core and cell sorting, the Department of Cardiothoracic Surgery at the University of Pittsburgh, in particular, Ms. Julie Ward for her help in coordination and gathering patient consents, as well as the Department of Cardiothoracic Surgery at the University of Colorado and the University of Colorado SPORE for providing samples, the Department of Hematology/Oncology at the University of Pittsburgh for research support and patient samples including C. Sander and M. Serrano, the Department of Otolaryngology at the University of Pittsburgh for providing patient samples and clinical information. This work was supported by the National Institutes of Health (R01 CA203689 and P01 AI108545 to D.A.A.V.), NCI Comprehensive Cancer Center Support CORE grant (CA047904 to D.A.A.V.), HNSCC SPORE grant (P50 CA097190 to R.L.F. and D.A.A.V), Melanoma SPORE grant and developmental research project (P50 CA121973 to J.M.K.), UPP Department of Surgery Foundation grant (internal grant to T.C.B.), NCI Institutional National Research Service Award in Cancer Therapeutics (T32 CA193205-04 to E. Chu) and an Eden Hall pilot grant (internal grant to A.S. and D.A.A.V.). This project also used the Hillman Cancer Center Immunologic Monitoring and Cellular Products Laboratory that is supported in part by award P30 CA047904.

## AUTHOR CONTRIBUTIONS

D.A.A.V and T.C.B. conceived and directed the project; D.A.A.V obtained funding for the project; A.S., T.C.B., and D.A.A.V conceptualized, designed, analyzed the experiments and wrote the manuscript; A.S. performed the experiments with help from A.R.C. for the single-cell RNAseq experiments and analyses; C. L, L.O., and M.V. obtained and processed multiple fresh and frozen samples from our patient cohorts; S.J. performed the qPCR for the TREC experiments; M.J.C. and S.C.W. performed and analyzed our STED microscopy; D.P.N performed the statistical analyses for our experiments; E.J.L., J.G.H, J.M.K, and R.L.F. obtained HD smoker control samples, NSCLC, metastatic melanoma, and HNSCC specimens and cohorts. All authors provided feedback and approved the manuscript.

## AUTHOR INFORMATION

The authors declare competing financial interests. D.A.A.V has submitted patents covering LAG3 and NRP1 that are licensed or pending and is entitled to a share in net income generated from licensing of these patent rights for commercial development. R.L.F. is on the advisory board and receives clinical trial and research funding from Bristol-Myers Squibb. E.L. is a consultant and receives research funding from Bristol-Myers Squibb.

## METHODS

### Human T cell populations

#### Patients and specimens

Patients were seen in the Hillman Cancer Center (HCC) at the University of Pittsburgh Medical Center (UPMC) or the Sidney Kimmel Comprehensive Cancer Center (SKCCC) and diagnosed with non-small cell lung carcinoma (NSCLC), head and neck squamous cell carcinoma (HNSCC), or metastatic melanoma. Patients were offered the option to donate blood or tissue as a part of our HCC research protocol (IRB# PRO16070383) or their corresponding SKCCC protocol. Patients then signed an IRB-approved informed consent and blood samples were obtained before initiation of treatment, but after pathologic diagnosis was established. Our fresh HNSCC cohort utilized for single-cell RNA sequencing (scRNAseq) included 11 tumor samples and 26 matched PBL and 6 HD PBL samples. The second cohort of fresh treatment-naïve HNSCC patients (n=49) was utilized for flow cytometric analysis and functional assays when sufficient cell yields allowed. The third cohort of banked PBL samples were thawed from HNSCC patients with both early-stage (n=24) and R/M (n=25) disease. Another cohort included fresh treatment-naïve NSCLC patients of all stages (n=50) utilized for flow cytometric analysis and functional assays when sufficient cell yields allowed. Our next cohort of banked PBL samples were thawed from NSCLC patients (n=33) along with matched HD smoker control PBL (n=10) utilized for flow cytometric analysis. Our fresh melanoma PBL cohort utilized 28 PBL samples for flow cytometric analysis and functional assays. An additional cohort of patients with advanced melanoma or other skin cancers provided PBL samples before checkpoint blockade and 12 weeks on checkpoint blockade (n=37) to the Sidney Kimmel Comprehensive Cancer Center at Johns Hopkins University.

#### Isolation of patient blood and tumor samples

Peripheral blood was drawn into both heparinized (flow cytometry and functional experiments) and EDTA tubes (for single-cell RNAseq experiments). PBL were isolated from peripheral blood by density gradient centrifugation using Ficoll-Hypaque (GE Healthcare Bioscience). Tumor samples were washed in RPMI containing antibiotics such as amphotericin B and penicillin-streptomycin for 30 minutes followed by mechanical and enzymatic digestion and further passage via a 100µm filter. Isolated PBL and TIL were then washed in RPMI media twice and utilized for downstream applications.

### Antibodies, Flow Cytometry, and STED microscopy

Single cell suspensions were stained with antibodies against CD45 (H130; BDbiosciences), CD45RA (H1100; Biolegend), CD4 (RPA-T4; Biolegend), CD8 (RPA-T8; Biolegend), surface LAG3 (1408; BMS), CD3 (SP34-2; BDbiosciences), ADAM10 (SHM14; Biolegend), PD1 (eBioJ105; eBioscience), ADAM17 (111633; R&D), pAKT (SDRNR; eBioscience), Ki67 (Ki67; Biolegend), total LAG3 (3DS223H; BMS), FoxP3 (PCH101, eBioscience), CCR7 (G043H7; Biolegend), CD62L (DREG56; Biolegend), CD103 (CD103; BDFisher), CD69 (FN50; Biolegend), CD95 (DX2; Biolegend), CD38 (HB-7; Biolegend), HLA-DR (L243; Biolegend), TNFα (MAb11; Biolegend), IFNγ (4S.B3; Biolegend), IL2 (MQ1-17H12; eBiosciences), granzyme-B (GB11; Biolegend), STAT1 Phospho (Ser727; Biolegend), STAT3 Phospho (Tyr705, 13A3-1; Biolegend), Bcl2 (100; Biolegend), TIGIT (MBSA43; eBiosciences), TIM3 (F38-2E2; Biolegend), surface NRP1 (12C2; Biolegend), intracellular NRP1 (nrp1; abcam), CD39 (A1; Biolegend). Dead cells were discriminated by staining with Fixable Viability Dye (eBioscience) in PBS. Cytokine expression analysis was conducted after a 4-hour activation of cells with PMA (0.1ng/mL; Sigma), Ionomycin (0.5ng/mL; Sigma), and Golgistop (Protein Transport Inhibitor; BDbioscience). Intracellular staining of cytokines and other proteins and transcription factors were conducted after fixation and permeabilization (eBioscience 88-8824-00) for 1 hour followed by two wash steps with permeabilization buffer (eBioscience). Surface and intracellular staining were conducted for 30 minutes at 4°C. Cells were sorted on MoFlo Astrios (Beckman Coulter). Flow cytometry experiments were performed using a Fortessa II (BD Bioscience). Flow cytometric data analyses were performed on FlowJo (Tree Star). Images were acquired with a Leica STED microscope after cell staining and fixation.

### Single-Cell RNA sequencing

#### Generation of single-cell RNAseq libraries

Libraries were prepared as previously described (Cillo et al, Immunity 2020). Briefly, live CD45^+^ cells (i.e. all immune cells) were sorted from PBL. Single-cell libraries were generated utilizing the chromium single-cell 3’ Reagent from V2 chemistry, as previously described^56^. Briefly, sorted cells were resuspended in PBS (0.04% BSA; Sigma) and then loaded into the 10X Controller for droplet generation, targeting recovery of 2,000 cells per sample. Cells were then lysed and reverse transcription was performed within the droplets and cDNA was isolated and amplified in bulk with 12 cycles of PCR. Amplified libraries were then size-selected utilizing SPRIselect beads, and adapters were ligated followed by sample indices. After another round of SPRIselect purification, a KAPA DNA Quantification PCR determined the concentration of libraries.

#### Sequencing of single-cell libraries

Libraries were diluted to 2nM and pooled for sequencing by NextSeq500/550 high-output v2 kits (at the Health Sciences Sequencing at the UPMC Children’s Hospital of Pittsburgh) for 150 cycles (parameters: Read 1: 26 cycles; i7 index: 8 cycles, Read 2: 98 cycles).

#### Demultiplexing, alignment, and generation of gene/barcode matrices

Raw sequence data were processed via CellRanger (10X genomics) and aligned to GRCh38 to generate feature barcode matrices. Cell barcodes with fewer than 3 UMI counts in 1% of cells and detection of 1% of cells were removed.

#### Alignment of data

Feature barcode matrices were read into Seurat v3, and data were integrated using a reciprocal principal component analysis (PCA) (T Stuart and A Butler et al, Cell 2019). Briefly, cells from each sample were log-normalized for library size, and highly variable features were identified from each sample. Features commonly expressed across samples were selected to integrate data. Next, data from each sample was scaled and PCA was performed based on highly variable genes. Reciprocal PCA was then performed using one sample each from a healthy donor, and HPV– HNSCC patient, and an HPV+ HNSCC patient to form an integrated dataset.

#### Visualization of all cells and identification of cell types

After creating an integrated dataset, we next utilized uniform manifold approximation and projection (UMAP; Becht E et al, Nature Biotech 2018) to visualize cells from all samples in a low-dimensional embedding. Cell types were identified as described previously (Cillo et al, Immunity 2020).

### Multivariate survival analysis of HNSCC cohort

Overall survival was defined as the time from baseline PBL sample acquisition until death from any cause. Between group-comparisons were completed via a log-rank test and multivariate analysis that included: stage, treatment course, race, gender, age group, tobacco use, alcohol use, histology, disease site (larynx, hypopharynx, oropharynx), and p16 status. The Kaplan-Meier method was utilized to calculate survival and their 95% confidence intervals.

### TREC qPCR assay

Samples and controls from fresh PBL were sorted by CD45RA^+^CCR7^+^CD62L^+^ cells from high LAG3^IC^ CD8^+^ T cell patients, low LAG3^IC^ CD8^+^ T cell patients, matched CD4^+^ T cells, HD CD8^+^ T cells, and effector cells from matched patients and donors. The patients included five healthy donors, three NSCLC patients, three metastatic melanoma patients, and four HNSCC patients with high LAG3 on their CD8^+^ T cells along with three NSCLC, three metastatic melanoma, and four HNSCC patients with low LAG3 on their CD8^+^ T cells. These samples were from our fresh sample accrual. Genomic DNA was extracted from cell pellets and then resuspended in nuclease-free water. Real-time qPCR settings were utilized per the MyTRECkit (genenplus) and run for 40 cycles with ROX master mix along with the built-in beta-actin and sample controls^38^. Samples were run in triplicate and data were analyzed after normalization of measurements.

### ATACseq

All libraries were sequenced with 75 bp paired-end reads. For each sample, we trimmed the nextera adapter sequences using cutadapt, and the resulting sequences were aligned to hg38 reference genome using bwa-mem. We filtered reads mapping to mitochondrial DNA from the analysis together with low-quality reads and reads in ENCODE blacklist regions. The model-based analysis of MACS2 was used to identify the peak regions with options “-q 0.01 -no-model -g hs”, and peak annotation was performed using HOMER.

### Functional Assays

#### Proliferation/stimulation assays

Bulk and naïve PBL were sorted on a MoFlo Astrios (Beckman Coulter) for CD8^+^ T cells, CD4^+^ T cells, and antigen-presenting cells (APCs). Autologous stimulation assays were performed with T cells, APCs, anti-CD3 (eBioscience, 0.5 µg/ml), and checkpoint blocking antibodies (anti-PD1 and anti-LAG3, 10ug/mL). T cells were analyzed for proliferation at day 5 via Cell trace dye (eBioscience) and inhibitory receptor expression (LAG3 and PD1) on a Fortessa II (BD Bioscience) and data analyses were performed via FlowJo (Tree Star).

#### Soluble Cytokine Analysis with Luminex

Plasma samples from patient cohorts were isolated by centrifugation and analyzed by ProcartaPlex 45 cytokine immunoassay (Thermo Fisher).

#### Inhibitory receptor induction assay via cytokine/plasma

LAG3^IC^ and PD1^IC^ expression were analyzed on bulk and naive T cells after incubation with various cytokines and patient plasma samples. Bulk and naïve CD8^+^ T cells and CD4^+^ T cells were sorted and incubated with low-dose IL2 (1000IU/mL), and low dose IL7 (1ng/mL) and no TCR stimulation. These assays included incubation with 45 individual cytokines at their corresponding EC_50_ concentration (R&D Systems) as well as plasma samples from our fresh patient PBL cohorts above with both low and high LAG3 CD8^+^ T cell subsets. Further assays included incubation with patient plasma and cytokine blockade including anti-IL6, anti-IL8, anti-IL22, anti-IL1b, anti-IL9, anti-IL10, anti-IL15, anti-IL21 at their corresponding EC_50_ concentration (Abcam). Incubation assays were conducted for 60 hours and analyzed by flow cytometry.

### LAG3 trafficking/shedding analysis

#### In vitro ADAM10 assays

Bulk and sorted peripheral CD8^+^ and CD4^+^ T cells were incubated in low dose IL2 and with and without varying concentrations of the ADAM10 and/or ADAM17 inhibitors: GI245023X (Tocris Biosciences), GW280264X (Aobious) inhibitors after 24 to 48 hours. The supernatant was collected and assayed for sLAG3 by ELISA in isolated cell types. Soluble LAG3 was also evaluated by ELISA in bulk and sorted peripheral CD8^+^ and CD4^+^ T cells *ex vivo* utilizing a sLAG3 ELISA kit (R&D Systems). A 96 well plate was coated with a capture antibody for 24 hours followed by sample and standard incubation. The samples were then washed with the corresponding primary, secondary HRP, and substrate solutions were added accordingly with wash steps in between. Absorbance was measured by an Epoch microplate spectrophotometer (Biotek Instruments). sLAG3 concentration was calculated using a purified sLAG3 standard curve, as previously described.^57^

### Statistical methods

Statistical analyses were conducted using Prism Version 7 (GraphPad). Comparisons of independent samples were conducted using Wilcoxon signed-rank tests (**Fig. 1c-g, 2b, c, 3a-f, 4a-h, Extended Fig. 1e, Extended Fig. 2a-i, 4a, b, e, f**). Analysis of nonlinear correlations were conducted utilizing a Spearman’s correlation coefficient (**Fig. 3c, 4c, d; Extended Fig. 2a-g, Extended Table 2**). No correlation designated as “nc”. Event-free survival was calculated with the Log-Rank (Mantel-Cox) test applied to Kaplan-Meier survival function estimates to determine statistical significance (**Fig. 1e**). “n” represents the number of human subjects used in an experiment, with the number of individual experiments listed in the legend. Samples are shown with the mean with or without error bars indicating standard error of the mean (s.e.m) *P <0.05, ***P <0.001, ****P< 0.0001, ns is P>0.05, Wilcoxon matched-pairs test.

