## Supplemental Figures for "Systemic immune dysfunction in cancer patients driven by IL6 and IL8 induction of an inhibitory receptor module in peripheral CD8^+^ T cells"

### Extended Figure 1: LAG3-dominant IR module is increased in peripheral CD8⁺ T cells in select patients by scRNAseq and flow cytometry

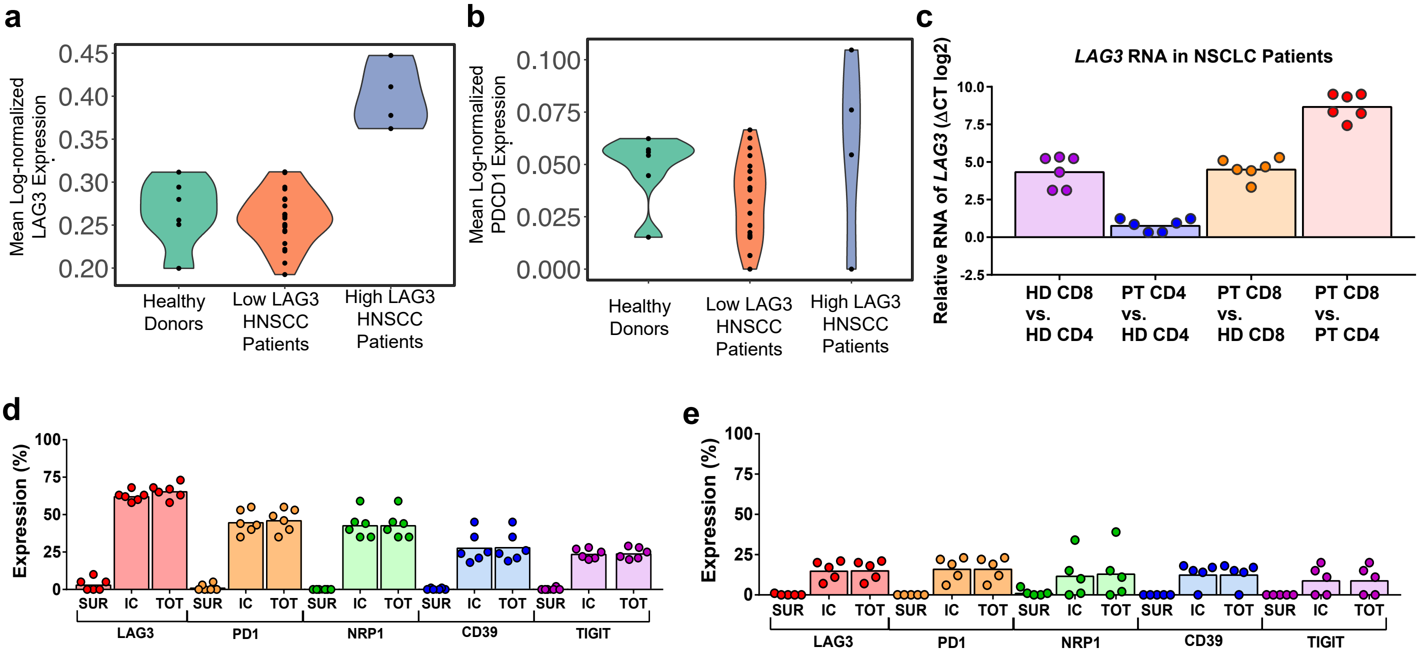


**Extended Figure 1.**

**a, b,** Mean log-normalized LAG3 and PDCD1 expression in CD8+ PBL from healthy donors (n=6), low LAG3 HNSCC patients (n=22), high LAG3 HNSCC patients (n=4).

**c,** Relative RNA expression of LAG3 (delta CT log2) by qPCR between healthy donor CD8+ T cells vs matched CD4+ T cells (purple; n=6), high $LAG3^{TOT}$ NSCLC patient CD4+ T cells vs. healthy donor CD4+ T cells (blue; n=6), high $LAG3^{TOT}$ NSCLC patient CD8+ T cells vs. healthy donor CD8+ T cells (orange; n=6), and high $LAG3^{TOT}$ NSCLC patient CD8+ T cells vs. matched patient CD4+ T cells (red; n=6).

**d, e,** Surface, intracellular and total LAG3 expression by flow cytometry of an total IR module consisting of LAG3, PD1, NRP1, CD39, TIGIT from only high total LAG3 patients (d) and low total LAG3 patients (e).

#### Extended Figure 2: LAG3-led IR module is increased in peripheral naïve CD8⁺ T cells in select patients by flow cytometry

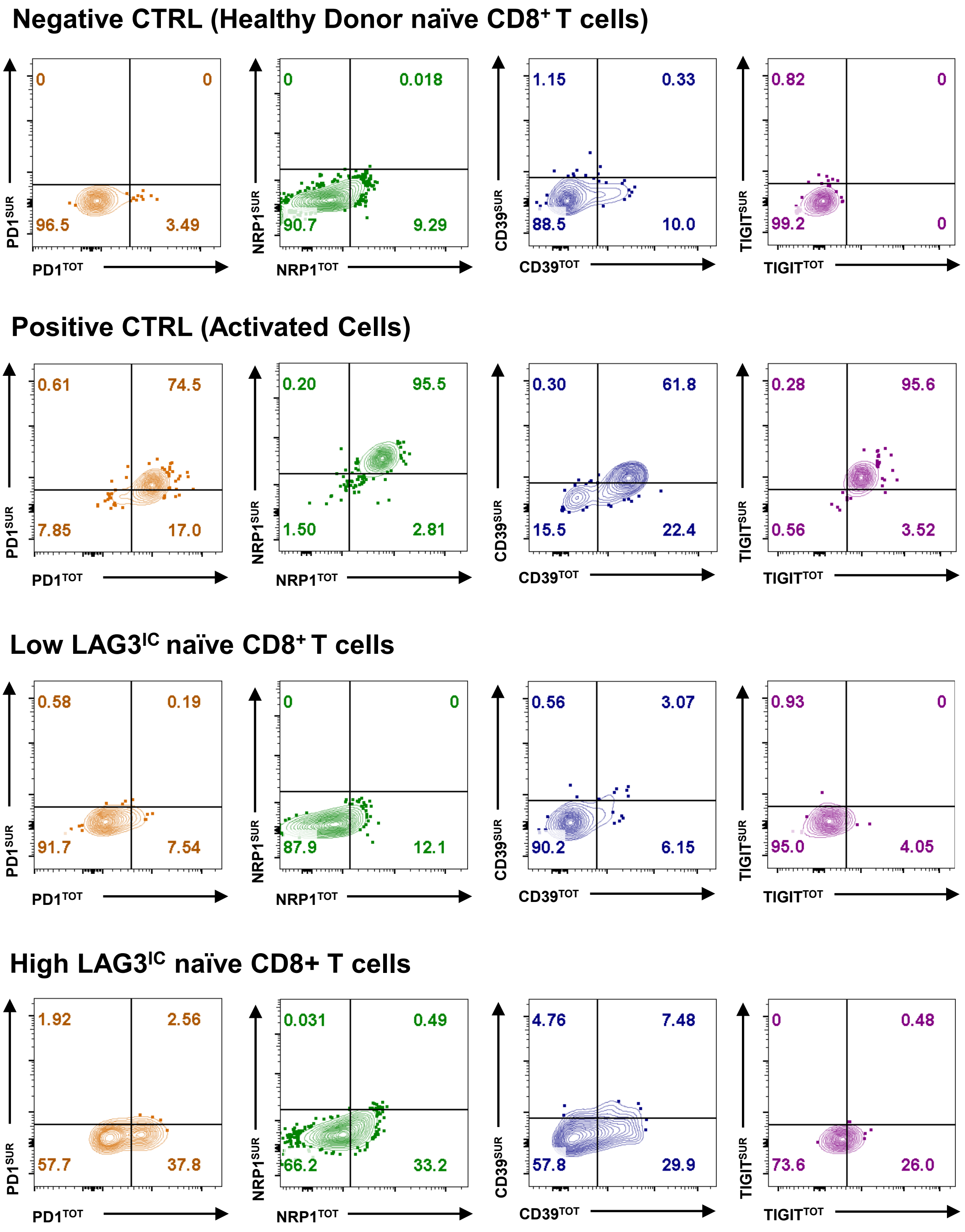


**Extended Figure 2.**
Representation flow cytometry plots of surface and intracellular (total) expression of PD1, NRP1, CD39, and TIGIT in the healthy donor CD8⁺ T cells (negative controls), activated CD8⁺ T cells (positive controls), low LAG3 CD8+ PBL and high LAG3 CD8+ PBL.

#### Extended Figure 3: Chromatin Accessibility of genes for IRs and T cell naivety

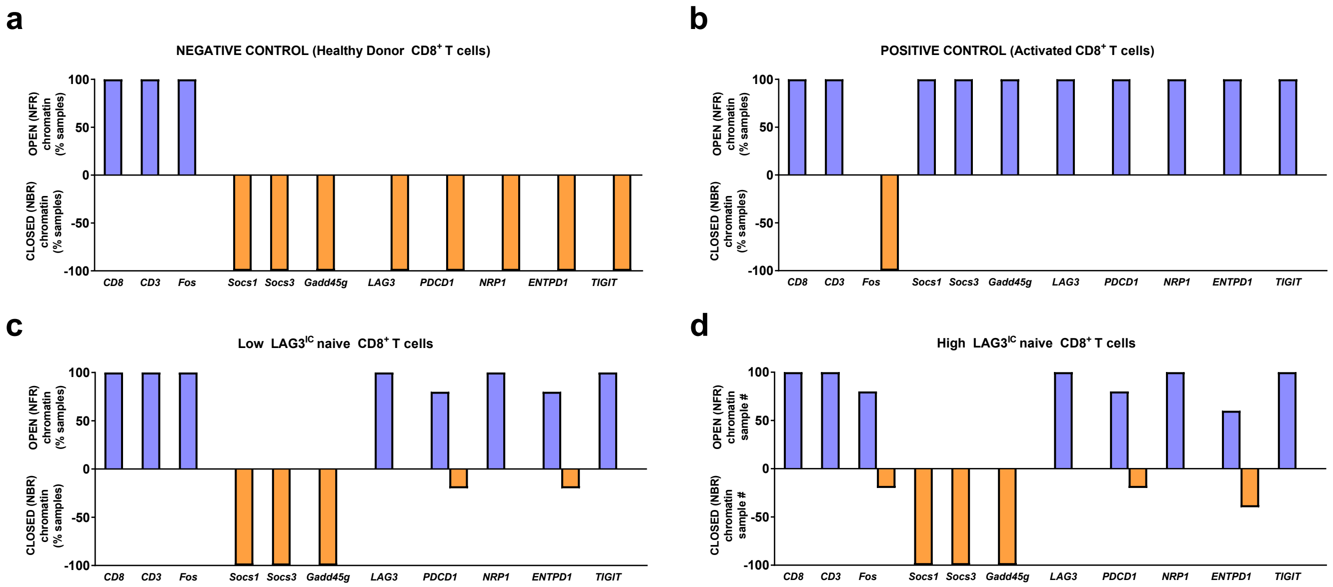


**Extended Figure 3.**
**a, b, c, d,** The percent of samples with “OPEN” chromatin or nucleosome free regions (NFR) at gene sites (blue) vs. “CLOSED” chromatin or nucleosome bound regions (NBR) at gene sites (orange) in healthy donors (n=2; negative control **a**), activated cells (n=2; positive control **b**), low LAG3 cells (n=5; **c**), high LAG3 cells (n=5; **d**). Evaluated chromatin sites for population markers as positive controls (*CD8, CD3*), sites for T cell naivete (*Fos*), and sites for activation (*Socs1, Socs3, Gadd45g*) and evaluation of sites of IR (*LAG3, PDCD1, NRP1, ENTPD1, TIGIT*).

#### Extended Figure 4: LAG3$^{IC}$ Expression by Ki-67 expression in CD8$^+$ T cells from responders and progressors on checkpoint blockade

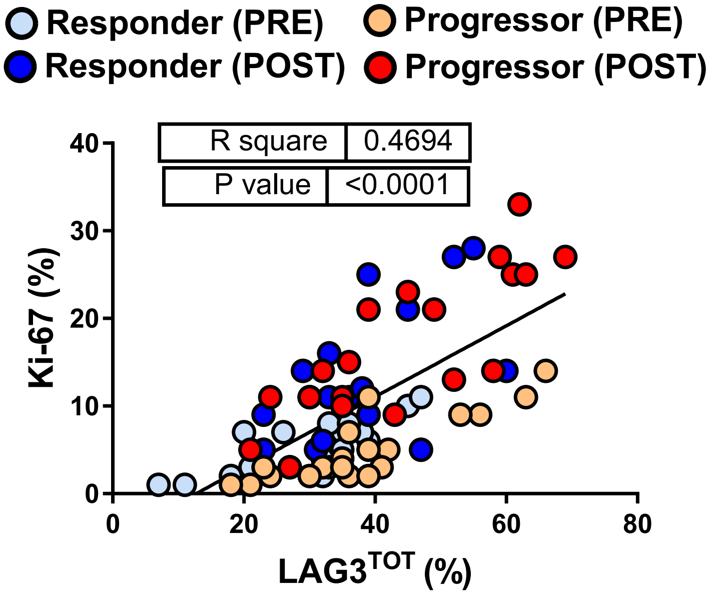


**Extended Figure 4.** Spearman's correlation of LAG3$^{IC}$ expression and ki-67 expression from peripheral CD8+ T cells from patients with advanced melanoma or skin cancers before (PRE) and 12 weeks after (POST) initial therapy.

#### Extended Figure 5: LAG3 is stored intracellularly in endosomal vesicles of CD8+ T cells

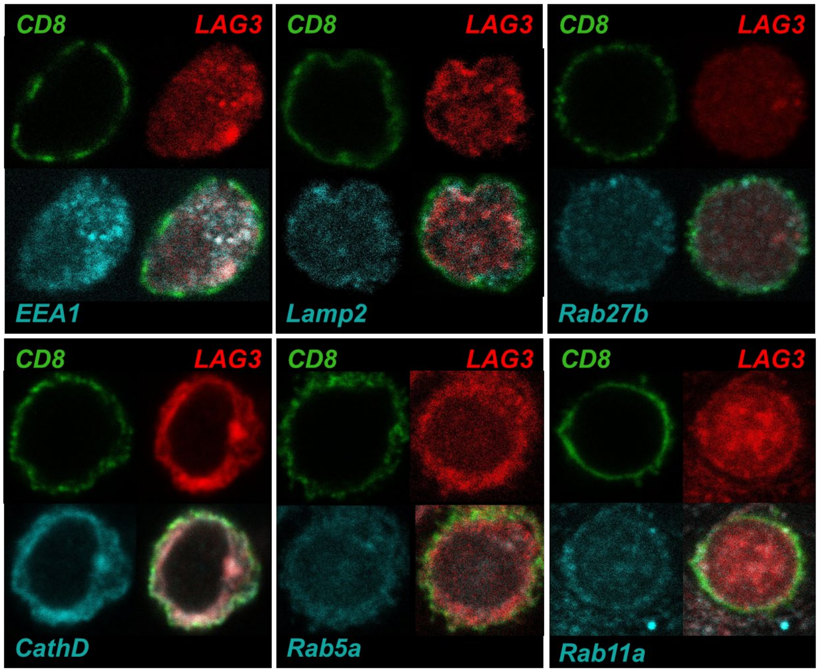


**Extended Figure 5.**
STED microscopy of LAG3 (red), CD8 (green), and markers of intracellular trafficking and storage (cyan): including: EEA1, Lamp2, Rab27b, Cathepsin D, Rab5a, Rab11a.

#### Extended Figure 6: Analysis of T cell Naivety in Peripheral CD8+ T cells can also express an elevated LAG3IC

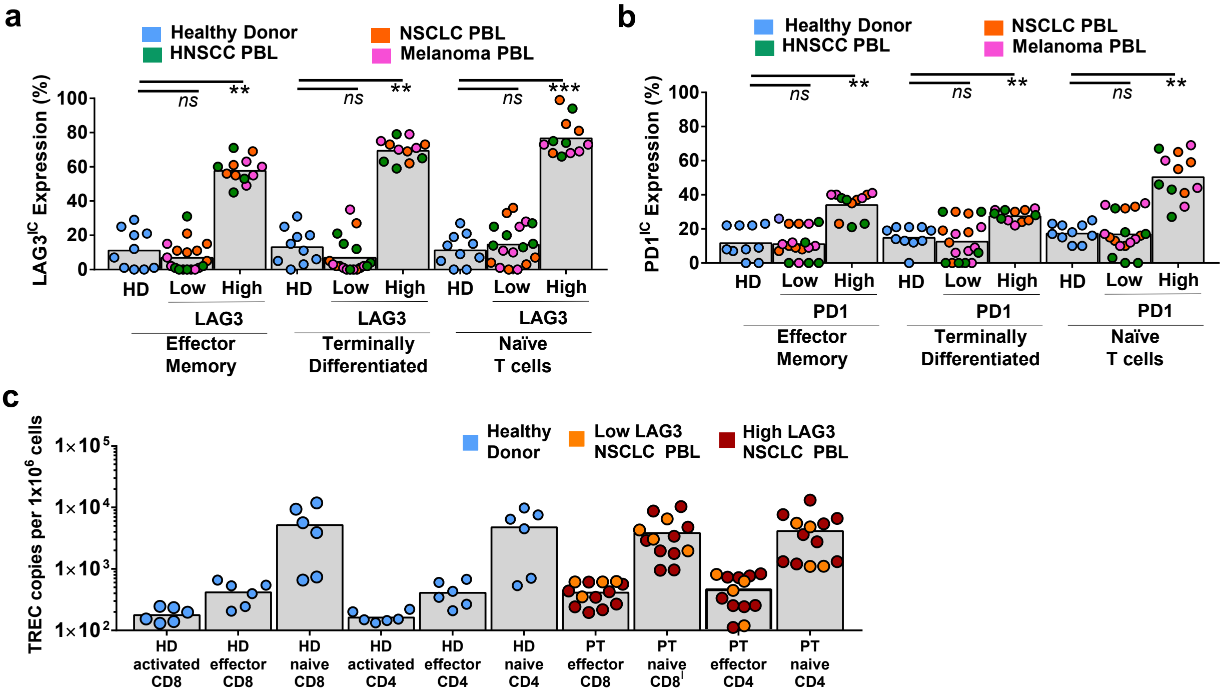


**Extended Figure 6.**
**a, b,** Analysis by flow cytometry of a prospective cohort of PBL (n=40) from patients with high LAG3/PD1 on $CD8^+$ T cells, low LAG3 on $CD8^+$ T cells and HD. LAG3/PD1 evaluated across T cells subsets including: naïve ($CD8^+$ $CD45RA^+$ $CCR7^+$ $CD62L^+$), effector memory ($CD8^+$ $CD45RA^-$ $CCR7^-$ $CD62L^-$), and terminally differentiated ($CD8^+$ $CD45RA^+$ $CCR7^-$ $CD62L^-$) T cells.[25]
**c,** TREC concentration as determined by qPCR of activated (positive control), effector, and naïve CD4+ and CD8+ T cells from healthy donors (n=6), low LAG3 NSCLC patients (n=4), and high LAG3 NSCLC patients (n=9). Statistical significance determined by Wilcoxon signed rank test (ns is P>0.05, * is P≤0.05, ** is P≤0.01, *** is P≤0.001, **** is P≤0.0001) comparing the change in proliferation after the addition of checkpoint blockade.

#### Extended Figure 7: Naïve Peripheral CD8$^{+}$ T cells can also express an elevated LAG3-dominant IR module leading to reduced function that can be partially rescued by anti-LAG3 or anti-PD1.

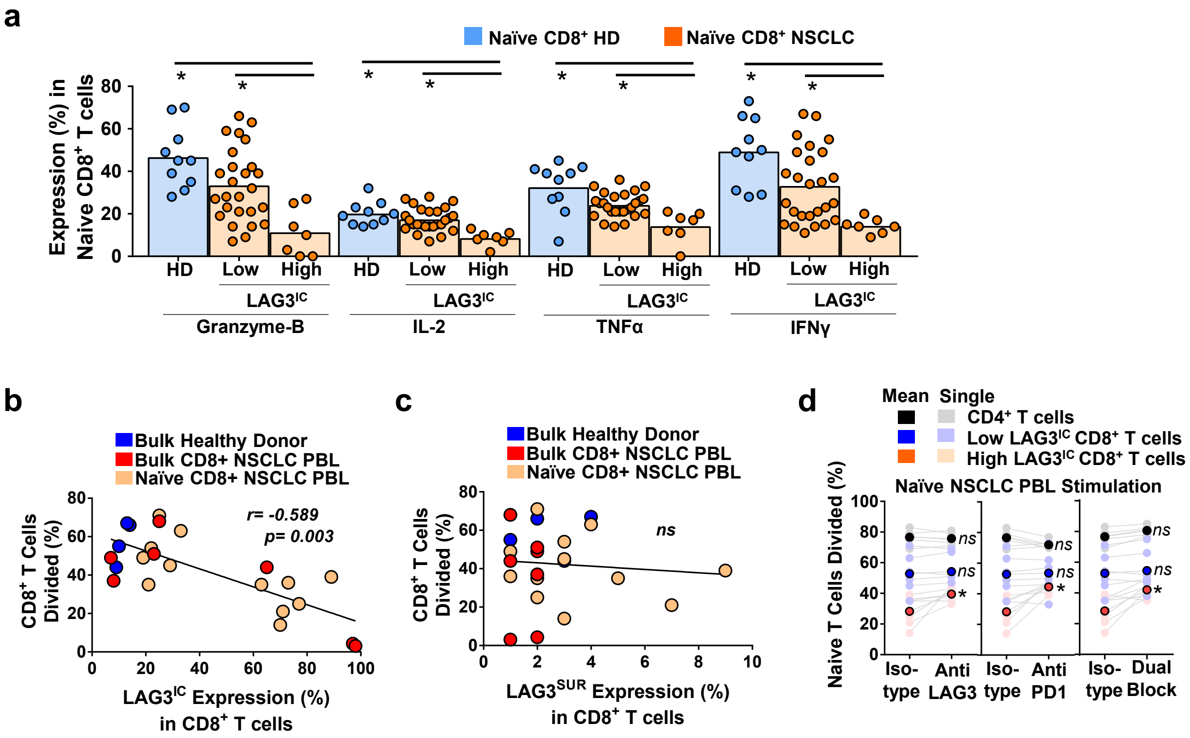


**Extended Figure 7.**

**a,** TNFα, IL2, IFNγ and granzyme B production evaluated by flow cytometry after 4-6 hour PMA and ionomycin stimulation in naïve (CD45RA$^{+}$CCR7$^{+}$CD62L$^{+}$) CD8$^{+}$ T cells with elevated LAG3$^{IC}$/PD1$^{IC}$ expression from NSCLC patients (n=7) compared to CD8$^{+}$ T cells with low LAG3$^{TOT}$/PD1$^{TOT}$ (n=26) and CD8$^{+}$ T cells from HD (n=10).

**b, c,** Spearman's correlation of proliferation and LAG3$^{IC}$ expression (e), or LAG3$^{SUR}$ expression (f) of patient bulk (n=7) and naïve (CD45RA$^{+}$CCR7$^{+}$CD62L$^{+}$; n=12) CD8$^{+}$ T cells subsets. Healthy donor (n=4) included in figure but not in correlation analysis.

**d,** naive CD4$^{+}$ T cells (n= 6 naïve) and CD8$^{+}$ T cells (n=11 naïve) from NSCLC patients with high (n=6 naive) and low LAG3$^{TOT}$ (n=5 naive) expression were stimulated with anti-CD3 (0.5µg/mL) and matched 1:1 APCs at a 96 hour timepoint in the presence of anti-LAG3 (10µg/mL) and/or anti-PD1 (10µg/mL). Statistical significance determined by Wilcoxon signed rank test (ns is $P>0.05$, * is $P\leq0.05$, ** is $P\leq0.01$, *** is $P\leq0.001$, **** is $P\leq0.0001$) comparing the change in proliferation after the addition of checkpoint blockade.

### Extended Figure 8: A LAG3-led IR module in naïve CD8⁺ PBL can be driven by IL6 or IL8 in the absence and presence of TCR stimulation.

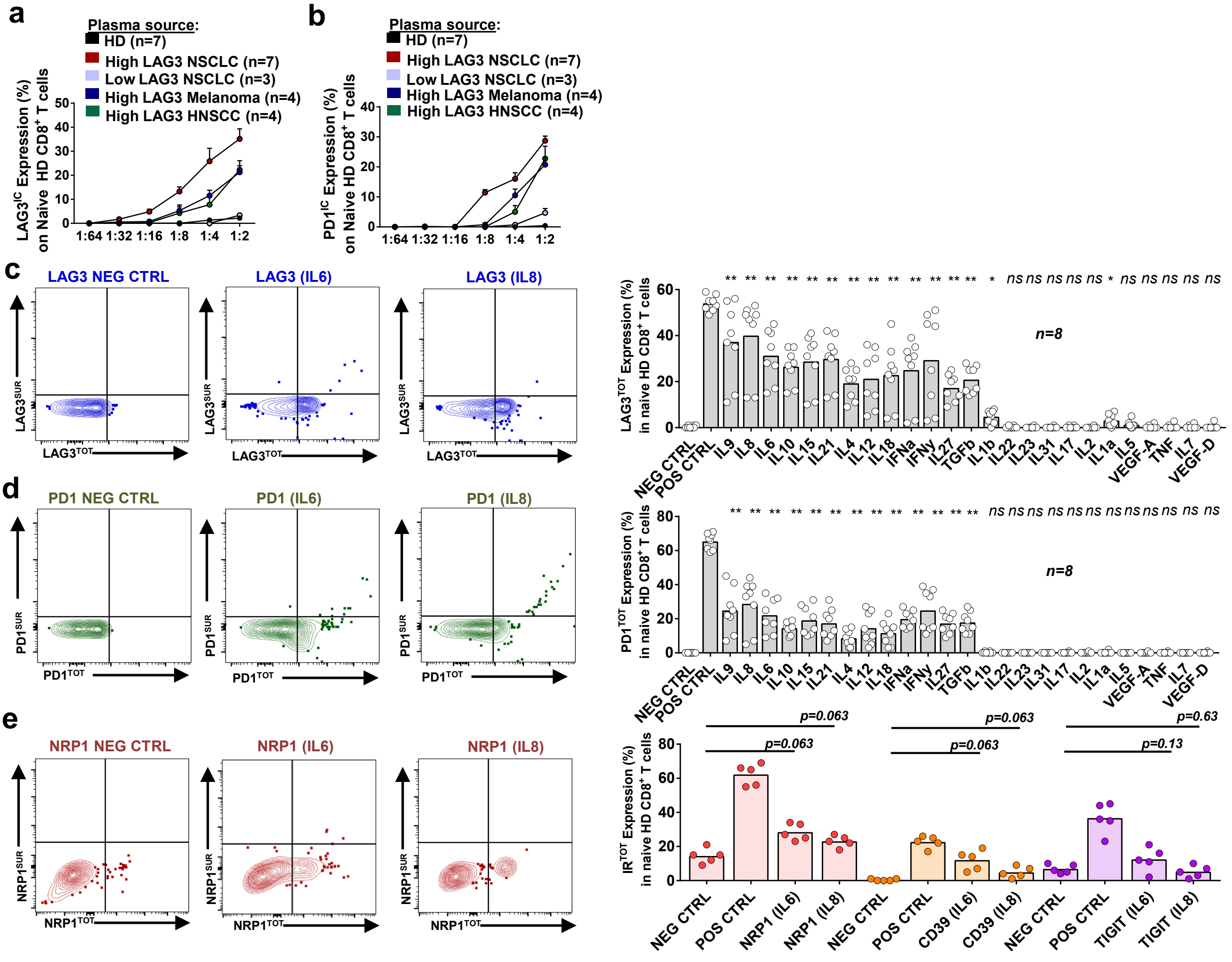


**Extended Figure 8.**

**a, b,** Naïve CD8⁺ HD T cells incubated with plasma from High LAG3$^{IC}$/PD1$^{IC}$ cancer patients in a similar manner to Fig 4. a, b. This plasma was known by multiplex to have elevated concentrations of all the cytokines of interest that correlated with LAG3$^{IC}$/PD1$^{IC}$. LAG3$^{IC}$ (a), and PD1$^{IC}$ (b), was evaluated by flow cytometry.

**c, d,** Representative flow plots and graphs of LAG3$^{TOT}$ expression (d) and PD1$^{TOT}$ expression (e) after a 72 hour incubation with naïve CD8+ PBL from healthy donors and a cytokine of interest at an EC50 concentration along with low dose IL2 and IL7. Negative control includes naïve CD8+ T cells with no additional cytokine and positive control are stimulated cells with TCR stimulation via anti-CD3, and anti-CD28.

**e,** NRP1$^{TOT}$, CD39$^{TOT}$, CTLA4$^{TOT}$, TIGIT$^{TOT}$ expression after 72 hour incubation with naïve CD8+ PBL from healthy donors and either IL6 or IL8 at an EC50 concentration along with low dose IL2 and IL7. Negative control includes naïve CD8+ T cells with no additional cytokine and positive control are CD8+ T cells that are stimulated with TCR stimulation via anti-CD3, and anti-CD28. Representative flow plots of NRP1 shown given trend as example.

Statistical significance determined by Wilcoxon signed rank test (ns is $P>0.05$, * is $P\leq0.05$, ** is $P\leq0.01$, *** is $P\leq0.001$, **** is $P\leq0.0001$). Error bars, s.d.

#### Extended Figure 9: A LAG3-dominant IR module in naïve $CD8^+$ PBL can be driven by IL6 or IL8 in the absence and presence of TCR stimulation.

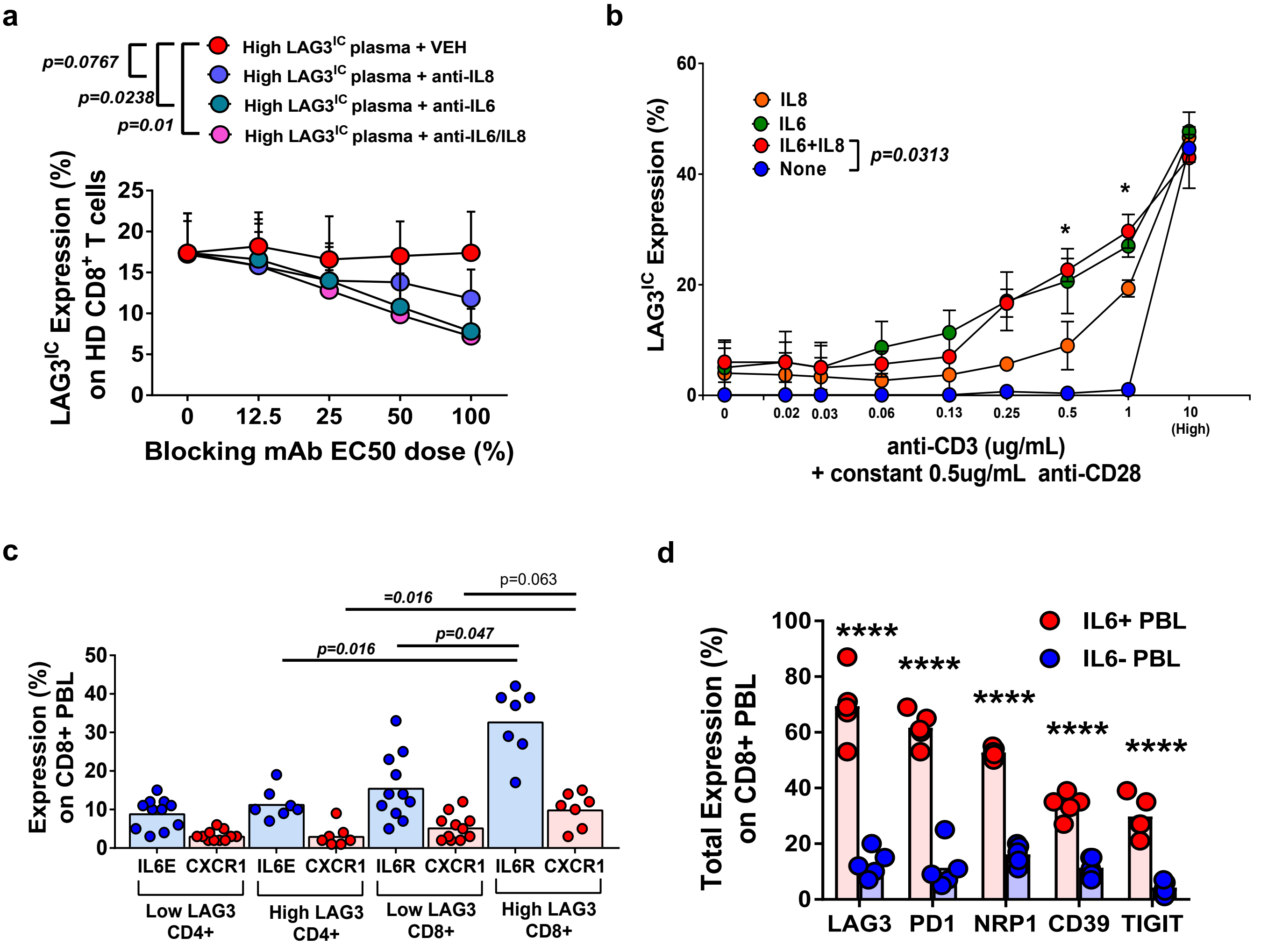


**Extended Figure 9.**

**a,** Naïve $CD8^+$ HD T cells incubated with plasma from High LAG3$^{IC}$/PD1$^{IC}$ metastatic melanoma patients in a similar manner to Fig 4. a, b. This plasma was known by multiplex to have elevated concentrations of all the cytokines of interest that correlated with LAG3$^{TOT}$/PD1$^{TOT}$. Incubation also included blockade of specific cytokines (anti-IL6, anti-IL8, anti-IL6/IL8, or vehicle) in an effort to rescue plasma induced LAG3$^{IC}$/PD1$^{IC}$expression. Concentrations of each blocking antibody varies, and the x-axis is noted as the percent of the reported $EC_{50}$ blocking dose.

**b,** LAG3$^{IC}$ expression after varying concentration of anti-CD3 and TCR stimulation for 72 hours with low dose IL2, and IL7, and anti-CD28 along with an EC50 concentration of IL8, IL6, or IL8+IL6, or no additional cytokine.

**c,** IL6 receptor (IL6R) expression and IL8 receptor (CXCR1) expression on CD8+ PBL and matched CD4+ PBL from NSCLC patients that have low LAG3 or high LAG3.

**d** Total expression of IRs including LAG3, PD1, NRP1, CD39, and TIGIT on CD8+ PBL from NSCLC patients that are either IL6+ or IL6-. Statistical significance determined by Wilcoxon signed rank test (ns is P>0.05, * is P≤0.05, ** is P≤0.01, *** is P≤0.001, **** is P≤0.0001). Error bars, s.d.

**Extended Table 1:**

| **Characteristic no. (%)** | **HNSCC (N=50)** |
| --- | --- |
| **Age-** | |
| >75 yr | 2 (4) |
| **Male sex** | 36 (72) |
| **Race-** | |
| White | 49 (98) |
| Asian | 0 (0) |
| Black | 1 (2) |
| **Stage-** | |
| Early | 25 (50) |
| Advanced | 25 (50) |
| **Tobacco use** | 44 (88) |
| **p16+** | 12 (24) |
| **Site-** | |
| Oropharynx | 5 (10) |
| Other | 45 (90) |

| **Characteristic** | **NSCLC (N=50)** |
| --- | --- |
| **Age-** | |
| >75 yr | 12 (24) |
| **Male sex** | 24 (48) |
| **Race-** | |
| White | 42 (84) |
| Asian | 2 (4) |
| Black | 6 (12) |
| **Stage-** | |
| Early | 42 (84) |
| Advanced | 8 (16) |
| **Smoker** | 47 (94) |
| **Histology-** | |
| Squamous | 14 (28) |
| Non-squamous | 36 (72) |
| +*EGFR* | 0 (0) |
| +*ALK* | 0 (0) |
| +*KRAS* | 0 (0) |

| **Characteristic** | **Melanoma (N=28)** |
| --- | --- |
| **Age-** | |
| >75 yr | 2 (7) |
| **Male sex** | 16 (57) |
| **Race-** | |
| White | 28 (100) |
| Asian | 0 (0) |
| Black | 0 (0) |
| **Stage-** | |
| Early | 22 (79) |
| Advanced | 6 (21) |
| ***BRAF*-** | |
| mutation+ | 12 (43) |
| mutation- | 16 (57) |

**Extended Table 1.**
Clinical characteristics of patient cohorts including HNSCC, NSCLC, and metastatic melanoma patients.

#### Extended Table 2: Advanced malignancies of the skin (JHU)

| | | Response (n=18) | Progression (n=19) |
|---|---|---|---|
| **Sex** | **Male** | 12 | 11 |
| | **Female** | 6 | 8 |
| **Age** | **>75yr** | 6 | 6 |
| | **<75yr** | 12 | 13 |
| **Histology** | **Melanoma** | 17 | 12 |
| | **Cutaneous SCC** | 1 | 0 |
| | **Basal Cell** | 0 | 2 |
| | **Mucosal Melanoma** | 0 | 3 |
| | **Uveal Melanoma** | 0 | 1 |
| | **Merkel Cell** | 0 | 1 |
| **Therapy** | **Pembro** | 6 | 5 |
| | **Nivo** | 8 | 6 |
| | **Nivo+Ipi** | 3 | 8 |
| | **Cemiplimab** | 1 | 0 |
| **Treatment History** | **Treatment-naïve** | 16 | 12 |
| | **Pretreated** | 2 | 7 |
| **Race** | **White** | 17 | 19 |
| | **Asian** | 0 | 0 |
| | **Black** | 1 | 0 |
| **BRAF-** | **Mutation +** | 6 | 4 |
| | **Mutation -** | 9 | 11 |
| | **Unknown** | 3 | 4 |
| **Stage** | **III** | 6 | 2 |
| | **IV** | 12 | 17 |
| **Response** | **CR** | 6 | N/A |
| | **PR** | 12 | N/A |
| | **PD** | N/A | 19 |

**Extended Table 2.**
Clinical characteristics of patient with advanced metastatic melanoma and other skin cancers from SKCCC.

#### Extended Table 3:

| Cytokine | Correlation with Inhibitory Receptor | LAG3$^{TOT}$ Induction (%) | PD1$^{TOT}$ Induction (%) |
|---|---|---|---|
| **IL9** | **(r=0.85)** | **37** | **25** |
| **IL8** | **(r=0.82)** | **40** | **29** |
| **IL6** | **(r=0.72)** | **31** | **22** |
| **IL10** | **(r=0.59)** | **27** | **15** |
| **IL15** | **(r=0.50)** | **29** | **19** |
| **IL21** | **(r=0.46)** | **30** | **17** |
| IL1b | **(r=0.74)** | 5.0 | 0.0 |
| IL22 | **(r=0.73)** | 0.0 | 0.0 |
| IL23 | **(r=0.73)** | 0.0 | 0.0 |
| IL31 | **(r=0.74)** | 0.0 | 0.0 |
| IL17 | **(r=0.65)** | 0.0 | 0.0 |
| IL2 | **(r=0.61)** | 0.0 | 0.0 |
| IL1α | **(r=0.61)** | 3.0 | 0.0 |
| IL5 | **(r=0.60)** | 1.0 | 0.0 |
| VEGF-A | **(r=0.57)** | 0.0 | 0.0 |
| TNFα | **(r=0.52)** | 0.0 | 0.0 |
| IL7 | **(r=0.52)** | 0.0 | 0.0 |
| VEGF-D | **(r=0.49)** | 0.0 | 0.0 |
| IL4 | (n.s.) | **19** | **9.0** |
| IL12 | (n.s.) | **21** | **14** |
| IL18 | (n.s.) | **23** | **12** |
| IFNα | (n.s.) | **25** | **20** |
| IFNγ | (n.s.) | **29** | **25** |
| IL27 | (n.s.) | **17** | **10** |
| TGFβ | (n.s.) | **21** | **19** |

**Extended Table 3:** Tabular list outlining cytokines and the correlation between their plasma concentrations and LAG3 expression on their naïve CD8$^{+}$ T cells followed by their ability to induce LAG3 expression in an *in vitro* incubation assay. Correlation was determined by Spearman's correlation noted as "rho" or "ρ" or as "nc" for "no correlation". $^{-}$ Cytokines that are at all correlated in plasma with LAG3$^{TOT}$ or PD1$^{TOT}$ expression on patient naïve CD8$^{+}$ T cells are noted in bold in the first column while the cytokines that are able to induce LAG$^{TOT}$/PD1$^{TOT}$ are noted in bold for the last two columns (see *Ext. Fig. 4e, f*). Cytokines that are both correlated and can induce LAG$^{TOT}$/PD1$^{TOT}$ expression are noted in red.
